# Recurrent neural network model of human event-related potentials in response to intensity oddball stimulation

**DOI:** 10.1101/2022.04.29.489982

**Authors:** Jamie A. O’Reilly

**Affiliations:** College of Biomedical Engineering, Rangsit University, Pathum Thani, Thailand

**Keywords:** Artificial neural networks, auditory neuroscience, computational neurophysiology, deviance detection, electrophysiology biomarkers, loudness dependence, predictive coding

## Abstract

The mismatch negativity (MMN) component of the human event-related potential (ERP) is frequently interpreted as a sensory prediction-error signal. However, there is ambiguity concerning the neurophysiology underlying hypothetical prediction and prediction-error signalling components, and whether these can be dissociated from overlapping obligatory components of the ERP that are sensitive to physical properties of sounds. In the present study, a hierarchical recurrent neural network (RNN) was fitted to ERP data from 38 subjects. After training the model to reproduce ERP waveforms evoked by 80 dB standard and 70 dB deviant stimuli, it was used to simulate a response to 90 dB deviant stimuli. Internal states of the RNN effectively combine to generate synthetic ERPs, where individual hidden units are loosely analogous to population-level sources. Model behaviour was characterised using principal component analysis of stimulus condition, layer, and individual unit responses. Hidden units were categorised according to their temporal response fields, and statistically significant differences among stimulus conditions were observed for amplitudes of units peaking in the 0 to 75 ms (P50), 75 to 125 ms (N1), and 250 to 400 ms (N3) latency ranges, surprisingly not including the measurement window of MMN. The model demonstrated opposite polarity changes in MMN amplitude produced by falling (70 dB) and rising (90 dB) intensity deviant stimuli, consistent with loudness dependence of sensory ERP components. Although perhaps less parsimoniously, these observations could be interpreted within the context of predictive coding theory, as examples of negative and positive prediction errors, respectively.

## 1. Introduction

Brains conserve energy expenditure associated with perception by efficiently modulating their responses to changing external environments. For example, cortical auditory-evoked responses normally decrease in magnitude with sound repetition, known as repetition-suppression, reflecting neurophysiological adaptations that optimize the efficiency of sound representations in accordance with their environmental significance (Harpaz et al., 2021; Weber and Fairhall, 2019). This description assumes that unreinforced repetitive sounds have downregulated environmental valence (Sara and Bouret, 2012), and lower magnitude cortical responses are more energy-efficient, following inhibitory homeostatic plasticity (Hertäg and Clopath, 2022). In contrast, responses to unexpected, novel sounds provoke comparatively enlarged cortical auditory-evoked potentials, reflecting a higher degree of environmental valence (Khouri and Nelken, 2015; Southwell and Chait, 2018). These findings are postulated to arise from a predictive model formed by the cortex that imposes top-down suppression of responses to expected afferent stimuli received from peripheral sensory organs, whereas unexpected stimuli trigger an enlarged bottom-up response that causes this predictive model to be updated, eliciting what is referred to as a sensory prediction-error signal (Garrido et al., 2009; Heilbron and Chait, 2018; Khouri and Nelken, 2015).

The preceding theory offers an explanation for mismatch negativity (MMN), a putative component of the human scalp-recorded event-related potential (ERP), which is widely thought to reflect sensory prediction-error signalling in response to unexpected environmental stimuli. This is usually observed experimentally in responses to auditory oddball stimuli. The MMN component is conventionally measured from the difference waveform between low-probability deviant and high-probability standard stimulus ERPs evoked by a passive oddball paradigm experiment (Näätänen et al., 2004). Changes in this signal are thought to reflect perceptual processing that can become impaired by underlying neuropathology, providing significant impetus for developing MMN as a clinical biomarker (Koshiyama et al., 2020; Schuelert et al., 2018; Taylor et al., 2017; Wang et al., 2022). However, considerable impediments to these efforts stem from ambiguity concerning the underlying neurophysiology of hypothetical prediction and prediction-error signalling components, and their dissociation from overlapping obligatory components that are sensitive to physical properties of sound (Heilbron and Chait, 2018; May, 2021; May and Tiitinen, 2010; O’Reilly and Conway, 2021; O’Reilly and O’Reilly, 2021). It has been suggested that MMN elicited by different physical changes in oddball stimuli may represent dissociable underlying neurophysiological processes (Näätänen et al., 2004; Nakajima et al., 2021; Pakarinen et al., 2007; Takegata et al., 2008; Todd et al., 2008), which perhaps implies a role for modulation of obligatory sensory components of the ERP by the physical properties of sounds in generating MMN (An et al., 2021; Hagenmuller et al., 2016; Kopp-Scheinpflug et al., 2018; O’Reilly, 2021a, 2021b; O’Reilly and Conway, 2021).

The present study follows up from (O’Reilly, 2021b), which found that 80 dB standard stimuli presented immediately after 70 dB deviant stimuli produce opposite polarity changes in ERP amplitudes compared with a 70 dB deviant following an 80 dB standard stimulus. Specifically, 70 dB stimuli produced more negative amplitude, whereas 80 dB stimuli that followed 70 dB stimuli produced more positive amplitude, within the measurement window of MMN, relative to consecutive 80 dB standard stimuli. This demonstrated that the magnitudes of auditory responses are influenced by immediately preceding sound intensity levels, leading to the hypothesis that louder deviant stimuli would similarly affect the MMN waveform. However, the dataset under consideration only included 80 dB standards and 70 dB deviants in an intensity-decrement oddball paradigm (Kappenman et al., 2021). The following article describes an attempt to circumvent this limitation and explore the intensity-dependence of ERP amplitudes during the latency range of MMN by developing a computational model using the existing data from 80 dB standard and 70 dB deviant stimuli and subsequently stimulating the response to an input signal representing a 90 dB deviant stimulus.

## 2. Material and methods

### 2.1 Data

Raw data used in this study were obtained from the ERP CORE (Compendium of Open Resources and Experiments), an open-access resource that includes normative data from several ERP experiments (Kappenman et al., 2021). The originating research was approved by the Institutional Review Board at the University of California, Davis, and all participants provided informed consent. Specifically, the ERP CORE MMN dataset was used, which contains EEG signals recorded from 40 healthy adult human subjects while they passively listened to an auditory oddball paradigm. The oddball sequence comprised 800 80 dB standard sounds and 200 70 dB deviant sounds. Both stimuli were 75 ms, 1000 Hz pure-tones with 5 ms rising and falling edge ramps. Raw EEG signals were band-pass filtered between 0.1 and 20 Hz and resampled to a sampling frequency of 200 Hz before re-referencing to the average of channels P9 and P10. Only data from channel FCz were modelled in the present study, which is the recommended site for analysing MMN in this dataset (Kappenman et al., 2021).

Standard stimuli that followed 70 dB deviant stimuli were excluded from analysis, as these were previously found to modulate the auditory response during the latency range of MMN (O’Reilly, 2021b). Fifteen initial standard trials were also ignored. From the remaining 585 standard trials, 200 were randomly selected in order to balance the numbers of standard and deviant trials. Subject 2 was removed because they were found to have fewer than 200 deviant trials, and subject 7 was removed because they were reported to display an excessive amount of artefacts (Kappenman et al., 2021). Electroencephalographic signals were segmented from −0.1 to 0.5 s about stimulus onsets, and a set of idealised trials for 80 dB standard and 70 dB deviant stimulus presentations were produced by averaging the trials from 38 subjects, producing an array of 200 idealised trials, each with 121 time-samples, for both stimulus conditions that were subsequently concatenated and used for model training.

### 2.2 Modelling

#### 2.2.1 Inputs

Input features consisted of step pulse functions designed to represent auditory stimuli with different amplitudes. Standard 80 dB stimuli were represented by a unit-step pulse that was high between 0 and 75 ms, whereas the input representing deviant 70 dB stimuli was half the amplitude of the standard input. Idealised trials produced by averaging data from the subjects in response to standard 80 dB (n = 200) and deviant 70 dB (n = 200) stimuli were shuffled together and provided as targets for the model to reproduce during training. Both input and output sequences were 121 samples in length, sampled at 200 Hz, representing −0.1 to 0.5 s about stimulus onset time. After training, the model response to a simulated 90 dB stimulus input, double the amplitude of the 80 dB standard, was produced. This modelling approach is illustrated in Figure 1a-c.

**Figure 1.**
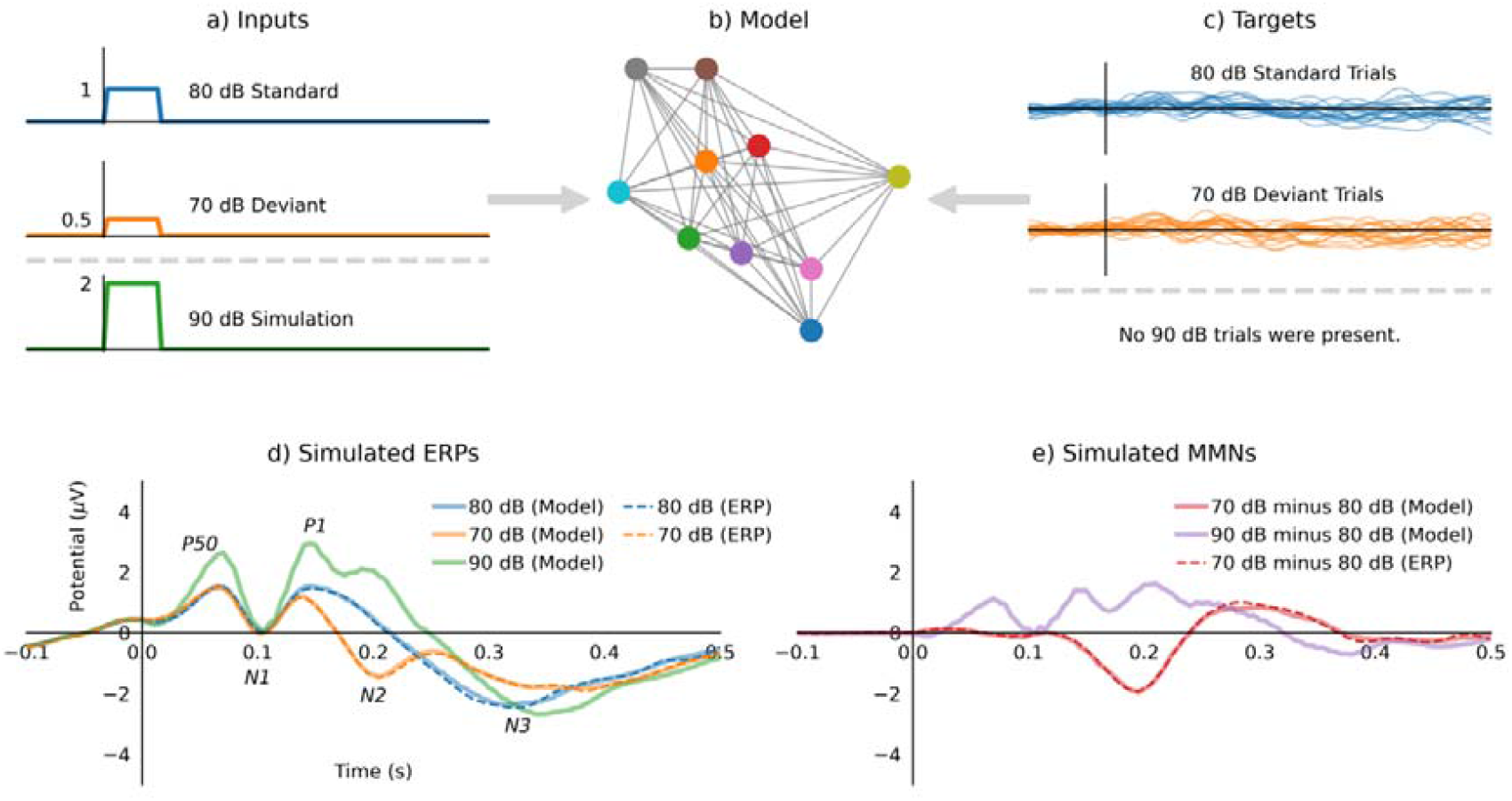
Hierarchical recurrent neural network modelling of human event-related potentials. a) Inputs to the model were amplitude modulated step pulse functions representing auditory stimuli. b) The model was an RNN with four hidden layers, each with 64 recurrent units and a single recurrent output unit. c) Targets presented to the model were idealised single-trials from stimuli presented in the intensity-decrement oddball paradigm. d) Model outputs after training, compared with grand-average event related potentials for 80 dB and 70 dB stimuli, including the simulated response to an input representing a 90 dB stimulus. e) Simulated mismatch negativity waveforms produced by subtracting the model 80 dB response from 70 dB and 90 dB responses. Latency ranges associated with peaks annotated in italics in (d) are defined in Table 1.

#### 2.2.2 Model architecture and training

The hierarchical recurrent neural network (RNN) model consisted of four hidden layers, each with 64 recurrent units, and a single recurrent output unit. Figure 1b is an illustrative, but not accurate, depiction of this model. The hidden units had rectified linear unit (relu) activation functions, while the output unit had a linear activation function. The model was trained for 1000 steps with a batch size of 100. During training, model parameters were adjusted to optimize mean squared error (MSE) difference between model outputs and targets using the back propagation through time (BPTT) algorithm implemented in Tensorflow (GoogleResearch, 2015). Adaptive moment estimation (adam) optimization was used with a learning rate of 0.001, beta-1 of 0.9, and beta-2 of 0.999. Forward connection weights were initialised from a Glorot uniform distribution (Glorot and Bengio, 2010), and recurrent connection weights were initialised from an orthogonal distribution (Saxe et al., 2013). This architecture was recently used to model mouse evoked epidural field potential data [*submitted for publication in Journal of Neural Engineering on 2 April 2022*].

#### 2.2.3 Validation

Five-fold cross validation was performed by randomly splitting idealised trials into 80% for training and 20% for validation. For each fold, the model was reinitialised and trained for 200 steps using 160 standard and 160 deviant trials, randomly shuffled together, with a batch size of 80. The training trials were averaged to produce training ERPs, and the 80 validation trials (40 standard and 40 deviant) were averaged to produce validation ERPs. After training each model, it outputs were compared with training and validation ERPs using Pearson’s correlation coefficient (r^2^). This produced overall r^2^ for training ERPs of 0.948, and for validation ERPs of 0.922 (all p < 0.0001), demonstrating sufficient capacity of the model to generalize the ERP data.

Leave one subject out (LOSO) cross-validation was also performed, where data from one subject were removed in turn and data from the remaining 37 subjects were averaged together to produce idealised trials used for training the model. Subsequently, model output was compared with training ERPs, produced by averaging the standard and deviant trials of the training data, and validation ERPs, produced by averaging the trials of the left-out subject. This process was repeated 38 times, each time training a reinitialised model for 200 steps, with a batch size of 100. Pearson’s correlation coefficients between model outputs and training and validation ERPs were recorded. The average overall r^2^ for the training data was 0.955 (all p < 0.0001), and for the validation data was 0.675 (p < 0.0001 for 69 out of 76 comparisons). A similar process was applied to grand-average ERP computation, where data from each subject were omitted in turn, producing an average r^2^ of 0.705 (p < 0.0001 for 71 out of 76 comparisons) between the left-out subject ERPs and grand-average ERPs. This demonstrates that the RNN effectively approximates the average of the target trials, equivalent to modelling the grand-average ERP waveform.

### 2.3 Model characterisation

#### 2.3.1 Simulated ERP and MMN waveforms

After training, the RNN was used to produce outputs representing grand-average ERP waveforms. The model outputs to input conditions representing 80 dB standards, 70 dB deviants, and simulated 90 dB deviants are plotted in Figure 1d, alongside grand-average ERPs for 80 dB and 70 dB stimuli for comparison. Difference waveforms were computed by subtracting the simulated 80 dB standard response from those of simulated 70 dB and 90 dB deviant responses, as shown in Figure 1e.

#### 2.3.2 Hidden unit activations

After training the RNN to generate ERP data, activities of its hidden units can be considered loosely analogous to biological sources that combine to produce genuine ERP waveforms. Drawing on this analogy, hidden units were analysed in an effort to derive insights concerning underlying computational principles that influence ERP and MMN waveform morphology. Matrix images displaying the activities of hidden units of each layer in response to 80 dB, 70 dB and 90 dB input conditions are plotted in Figure 2 along with respective model outputs.

**Figure 2.**
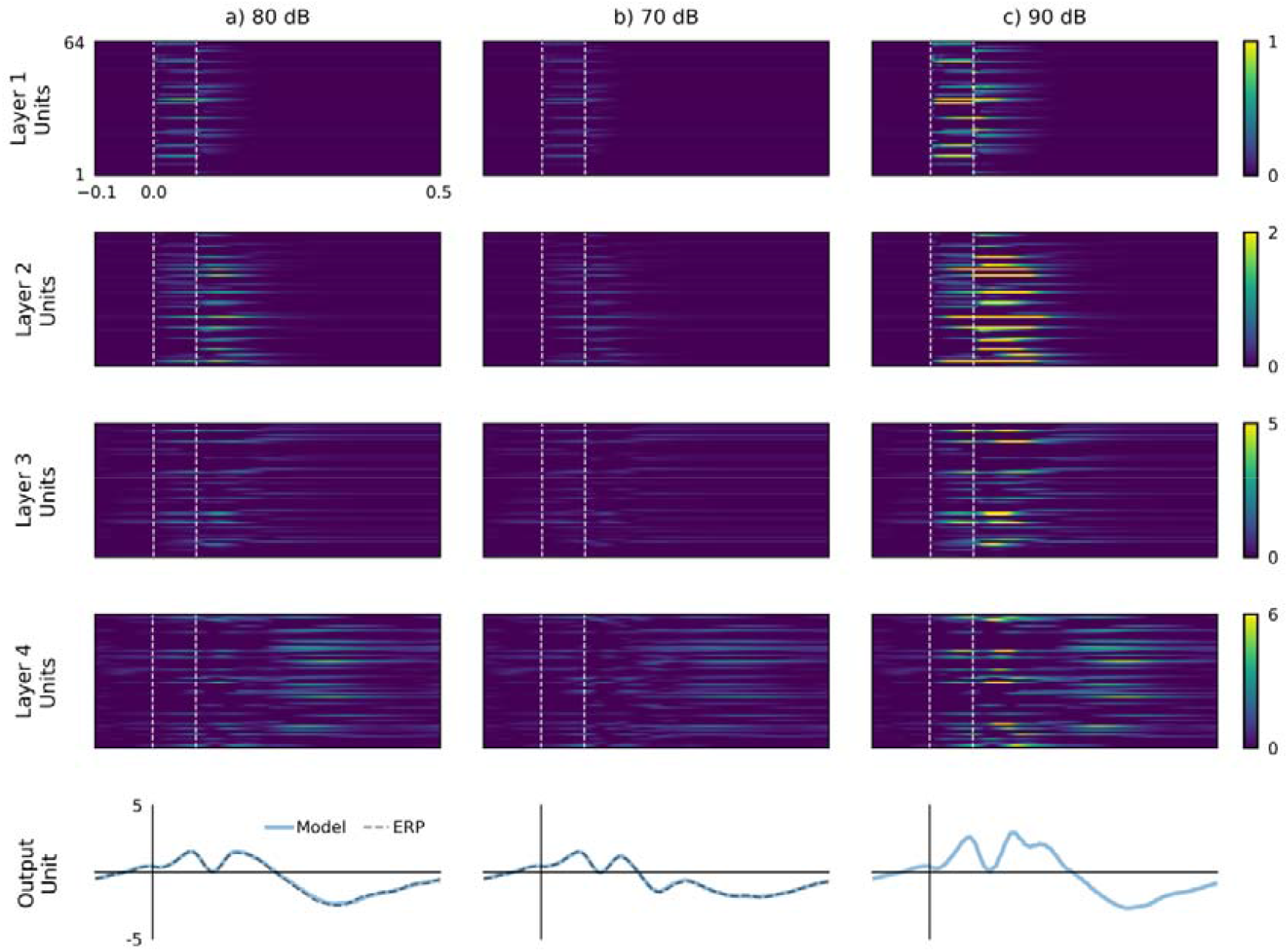
Model outputs and hidden layer activations plotted by layer and input condition. Responses to input representations of (a) 80 dB, (b) 70 dB, and (c) 90 dB stimuli are presented in columns. Amplitudes of hidden unit activations, shown in the four upper rows, increase towards the output. Patterns of hidden unit activations also spread out and become more complex with increasing layer depth. There are also evident differences in hidden unit response magnitudes between input conditions. Outputs plotted in the bottom row exhibit close agreement between 80 dB and 70 dB model outputs with their respective grand-average event related potentials. Vertical white dashed lines in the four upper rows represent the window during which stimulation was applied. All panels display data from −0.1 to 0.5 s about stimulus onset time at 0 s.

Histograms of hidden unit peak latencies in each layer were used to determine suitable boundaries for unit categorisation based on temporal response fields. These latency ranges also coincided with prominent deflections in the grand-average ERP waveforms: 0 to 75 ms (P1), 75 to 125 ms (N1), 125 to 175 ms (P2), 175 to 250 ms (N2), and 250 to 400 ms (N3). Some units peaked before stimulus onset, and some were silent. Numbers of units categorised according to these criteria are detailed in Table 1. Time-domain plots of hidden unit activations plotted by layer and input stimulus condition are also presented in Figure 3.

**Figure 3.**
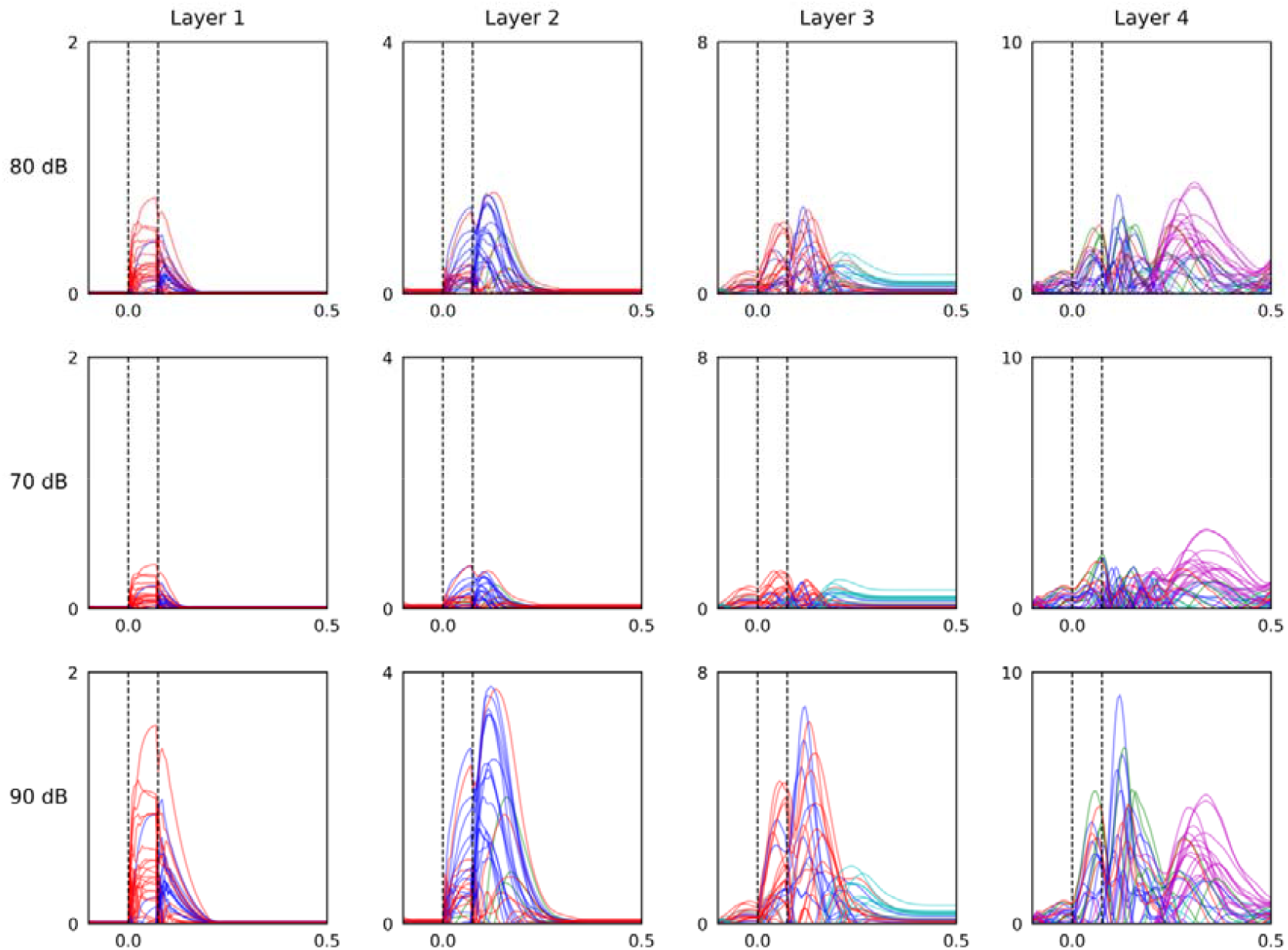
Time-domain analysis of hidden unit activations categorised by peak latency. Layers are plotted by column and stimulus conditions are plotted by row. Hidden units were categorised into seven classes based on their peak latency: silent (black), <0 ms (grey), 0 to 75 ms (red), 75 to 125 ms (blue), 125 to 175 ms (green), 175 to 250 ms (cyan), and 250 to 400 ms (magenta). This figure also illustrates differences in magnitude between hidden layer activations and model response to simulated 70 dB, 80 dB, and 90 dB stimulus conditions. Vertical black dashed lines represent stimulus-on and off-times, and data is plotted from −0.1 to 0.5 s about stimulus-onset.

**Table 1.** Model hidden unit categorisations by peak latency

| Peak | Latency range | Layer 1 | Layer 2 | Layer 3 | Layer 4 | Total |
| --- | --- | --- | --- | --- | --- | --- |
| Silent | No peak | 21 | 23 | 22 | 12 | 78 |
| Pre-stimulus | < 0 ms | 0 | 1 | 3 | 8 | 12 |
| P50 | 0 to 75 ms | 27 | 15 | 18 | 4 | 64 |
| N1 | 75 to 125 ms | 15 | 18 | 11 | 12 | 56 |
| P2 (MMN) | 125 to 175 ms | 1 | 6 | 0 | 6 | 13 |
| N2 (MMN) | 175 to 250 ms | 0 | 1 | 10 | 5 | 16 |
| N3 | 250 to 400 ms | 0 | 0 | 0 | 17 | 17 |
| **Total** | - | 64 | 64 | 64 | 64 | 256 |

The most frequent latency range for peak activity across input conditions was used to determine the category for each hidden unit.

#### 2.3.3 Principal component analysis

Model hidden unit activations were also characterised using PCA. Three different approaches were taken: (i) analysing model responses to three different stimulus conditions, (ii) analysing responses of different layers, and (iii) analysing individual units in each layer. In each case, data were reduced to two principal components (PC1 and PC2), as shown in Figure 4.

**Figure 4.**
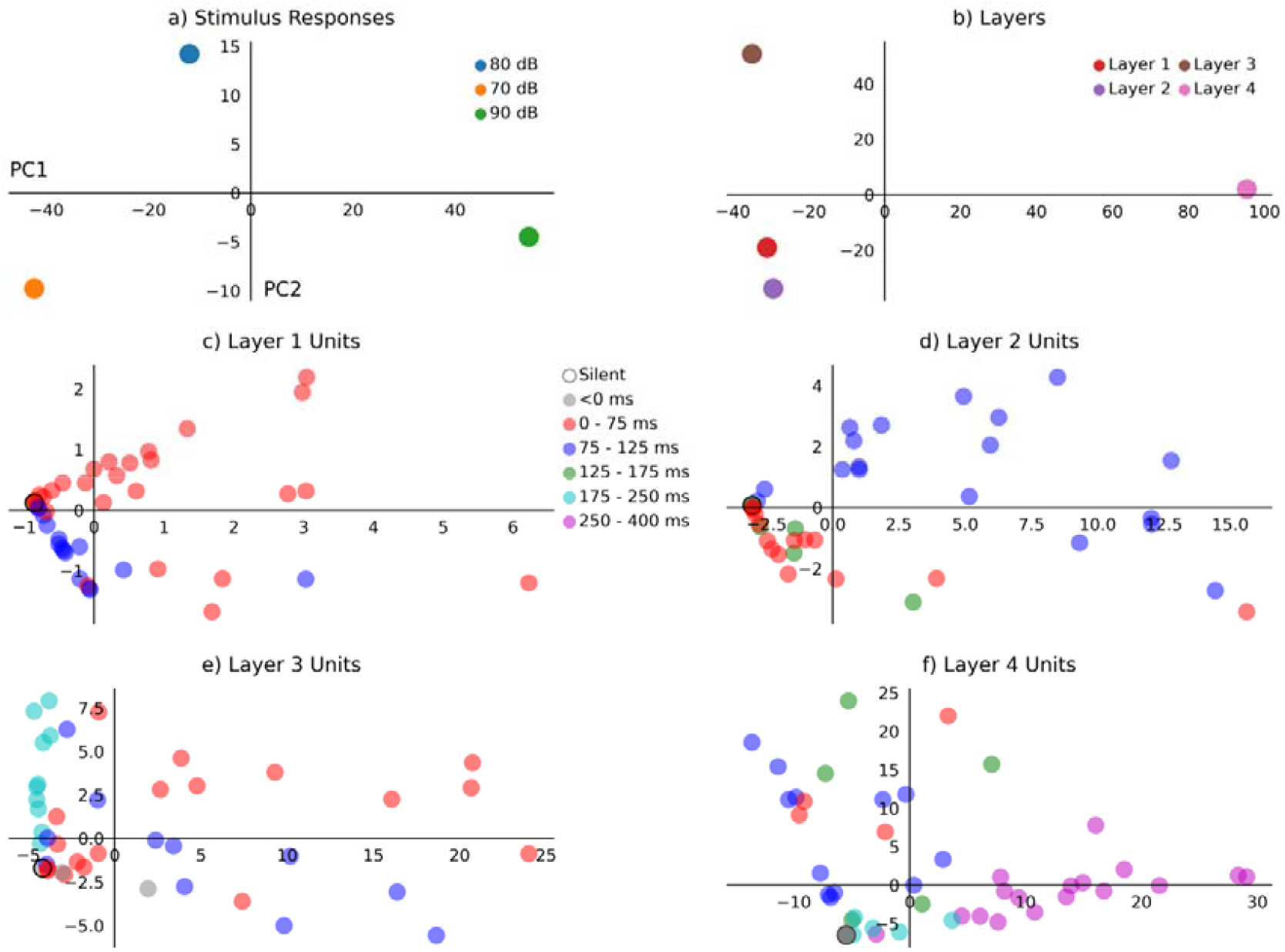
Principal component analysis of hidden unit activations characterises model behaviour. a) Responses to simulated input conditions show separation by intensity along the first principal component (PC1), and by standard or deviant identity along the second principal component (PC2). b) Analysis of layers indicates that layer 4 was the main source of variance in PC1, whereas layer 3 was more involved in PC2 variance. Principal components from individual hidden units are plotted in (c) to (f) for layers 1 to 4, with points coloured to represent unit categorisations described in Table 1, showing partial clustering of unit categories.

### 2.4 Statistical analysis

Pearson’s correlation coefficient (r^2^) was used to compare model outputs with grand-average ERP waveforms. One-way analysis of variance (ANOVA) tests were performed to compare hidden unit peak activations among stimulus conditions, and Bonferroni corrections for multiple comparisons were applied post-hoc. These tests were performed by unit category and by layers, as shown in Figure 5. Fifteen separate tests were performed hence p-values were adjusted to reflect a threshold for statistical significance of 0.05/15. Following a statistically significant finding from an ANOVA test, pairwise two-tailed dependent t-tests were performed between data evoked by each stimulus condition.

**Figure 5.**
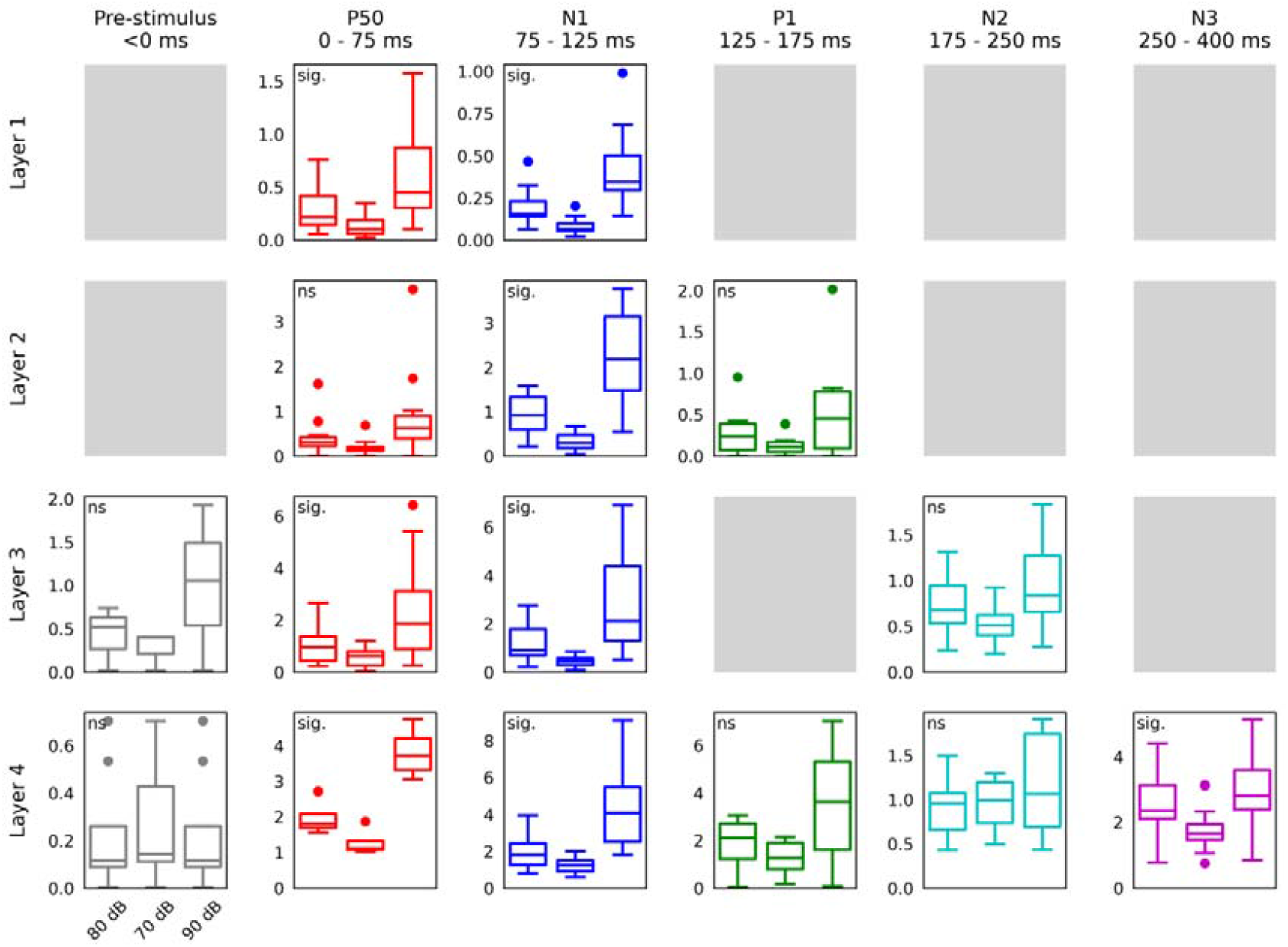
Analysis of hidden unit peak amplitudes by layer and peak latency range. Boxplots depict measurements in response to different stimulus conditions, and centrelines represent the median. Layer responses are presented row-wise, and unit categories are plotted column-wise. Layers with fewer than two units in a given category are greyed-out. Results of one-way ANOVA are annotated in the top-left corner of each panel as either not significant (ns) or statistically significant (sig.) based on p-values corrected for fifteen multiple comparisons using the Bonferroni adjustment (i.e. p < 0.05/15). Stimulus condition has a pronounced influence on P50 and N1 units across most layers, and also affects N3 units in layer 4.

### 2.5 Software

In this study, Python 3 was used with Matplotlib 3.3.3 (Hunter, 2007), MNE 0.23.4 (Gramfort et al., 2013), Numpy 1.19.5 (Harris et al., 2020), Scipy 1.7.1 (Jones et al., 2015), and Tensorflow 2.4.1 (GoogleResearch, 2015).

## 3. Results

The modelling approach and output waveforms are illustrated in Figure 1. After training the model, its outputs to simulated 80 dB standard and 70 dB deviant stimuli were nearly identical to grandaverage ERP waveforms, with r^2^ = 0.998 (p < 0.0001) and r^2^ = 0.999 (p < 0.0001), respectively. Likewise, the resulting simulated MMN waveform was also in close agreement with the human data, with r^2^ = 0.995 (p < 0.0001). Although there is no ground-truth waveform for comparison, the model output in response to a simulated 90 dB deviant input appears to follow a pattern consistent with intensity modulation. In the simulated MMN waveform produced by subtracting the model response to 80 dB input from that of 90 dB input, the amplitude difference observed across the latency of MMN was of opposite polarity to that produced by the 70 dB deviant.

Model hidden unit activations and outputs are plotted in Figure 2. This figure displays activations of 64 units from each hidden layer of the hierarchical RNN in image format, and output unit activation plotted as time-domain waveforms. Information flow though the network, represented by hidden unit activations, can be observed to spread out and increase in complexity from the first (shallowest) to the fourth (deepest) hidden layer. It is also evident that hidden units exhibit higher magnitude activations in response to louder simulated stimuli.

Hidden units were individually classified into seven different categories, as outlined in Table 1. Silent units did not display any activity above zero, of which 78/256 (30.4%) of hidden units were categorised, displaying a fair degree of sparsity or model redundancy. Pre-stimulus units were those whose activity peaked before stimulus onset, consisting of 12/256 (4.7%) of all hidden units, representing the trial overlap response observed in the pre-stimulus baseline. Units whose activity peaked during the 0 to 75 ms window were classed as P50 units and comprised 64/256 (25%) of the hidden units. The N1 units had activity that peaked within the 75 to 125 ms window, which included 56/256 (21.9%) of the hidden units. The P2 units peaked from 125 and 175 ms, and encompassed 13/256 (5.1%), while N2 units that peaked from 175 to 250 ms accounted for 16/256 (6.3%) of the model’s hidden units. Both P2 and N2 units peaked within the latency range of MMN, and if considered together account for 29/256 (11.3%) of the hidden units. The final window, from 250 to 400 ms, captured 17/256 (6.6%) of the hidden units that were associated with the slower N3 peak. Time-domain graphs of hidden unit activations are shown in Figure 3, with traces coloured according to the aforementioned unit categorisations. These waveforms indicate partial intensity dependence of some classes of units in different layers, and also display increasing temporal complexity of hidden unit activations in deeper layers.

To perform PCA of model responses to stimulus conditions, activations of hidden units were concatenated, producing a matrix of 3 (stimulus conditions) by 30976 (4 layers, 64 units, and 121 samples). This matrix was transformed into two principal components that explained 100% of the variance in the data. The results plotted in Figure 4a demonstrate separation of 70 dB, 80 dB, and 90 dB inputs by PC1, and separation of the 80 dB standard from 70 dB and 90 dB deviants by PC2. Similarly, PCA by layer was performed by reducing a matrix of 4 (layers) by 23232 (3 stimulus conditions, 64 units, 121 samples) into two components that accounted for 93.4% of its variance. Figure 4b shows that layer 4 and layer 3 responses are primarily responsible for the first and second principal modes of variance, respectively, whereas layer 1 and layer 2 responses are closer together in principal component space. Units in each layer were also treated individually in a comparable manner. These were transformed from 64 (units) by 323 (3 stimulus conditions, 121 samples) into two-dimensional principal component space. For layers 1, 2, 3, and 4, two principal components accounted for 96.4%, 94.2%, 86.3, and 80.1% of the variance in the data, respectively. The results of PCA applied to layers are shown in Figure 4c-f, which portray partial clustering of units by their temporal response category described above.

Hidden unit peak amplitude data is given by unit category and layer in Figure 5. One-way ANOVA tests were performed to compare the effects of stimulus condition on unit peak activations, with p-values adjusted using the Bonferroni correction for multiple tests. In layer 1, units peaking within the P50 latency range of 0 to 75 ms post stimulus showed a statistically significant effect of stimulus condition (F(2,24) = 22.1, p = 3.71e-07), with pairwise comparisons all demonstrating statistically significant differences (80 dB vs. 70 dB: t(25) = 7.91, p = 2.18e-08, 80 dB vs. 90 dB: t(25) = −7.95, p = 1.99e-08, and 70 dB vs. 90 dB: t(25) = −7.94, p = 2.05e-08). Layer 3 P50 units also showed a statistically significant effect (F(2,15) = 10.1, p = 0.003) between all conditions (80 dB vs. 70 dB: t(16) = 4.79, p = 0.00017, 80 dB vs. 90 dB: t(16) = −4.8, p = 0.000167, and 70 dB vs. 90 dB: t(16) = −4.82, p = 0.000161), as did layer 4 (F(2,1) = 21.2, p = 0.00593, 80 dB vs. 70 dB: t(2) = 8.88, p = 0.00301, 80 dB vs. 90 dB: t(2) = −11.9, p = 0.00129, and 70 dB vs. 90 dB: t(2) = −11.4, p = 0.00143). Units peaking in the N1 latency range of 75 to 125 ms also showed a statistically significant effect of stimulus condition in layer 1 (F(2,12) = 23.2, p = 2.5e-06, 80 dB vs. 70 dB: t(13) = 7.85, p = 1.72e-06, 80 dB vs. 90 dB: t(13) = −7.65, p = 2.29e-06, and 70 dB vs. 90 dB: t(13) = −7.72, p = 2.07e-06), layer 2 (F(2,15) = 37.9, p = 1.23e-09, 80 dB vs. 70 dB: t(16) = 9.4, p = 3.83e-08, 80 dB vs. 90 dB: t(16) = −9.11, p = 5.97e-08, and 70 dB vs. 90 dB: t(16) = −9.21, p = 5.11e-08), layer 3 (F(2,8) = 9.23, p = 0.0113, 80 dB vs. 70 dB: t(9) = 4.12, p = 0.00208, 80 dB vs. 90 dB: t(9) = −4.03, p = 0.00239, and 70 dB vs. 90 dB: t(9) = −4.06, p = 0.00227), and layer 4 (F(2,9) = 15, p = 0.000341, 80 dB vs. 70 dB: t(10) = 3.55, p = 0.00458, 80 dB vs. 90 dB: t(10) = −5.86, p = 0.000109, and 70 dB vs. 90 dB: t(10) = −5.14, p = 0.000322). Peak amplitudes from units in the P2 latency range of 125 to 175 ms, or N2 latency range of 175 to 250 ms. In layer 4, peak amplitudes of units in the N3 latency range between 250 and 400 ms were significantly affected by stimulus condition (F(2,14) = 8.1, p = 0.014), which was also determined to be due to input intensity differences (80 dB vs. 70 dB: t(15) = 9.3, p = 7.43e-08, 80 dB vs. 90 dB: t(15) = −9.55, p = 5.18e-08, 70 dB vs. 90 dB: t(15) = −9.44, p = 6.08e-08).

## 4. Discussion

Simulated ERPs from the model and resulting difference waveforms (Figure 1d-e) appear to support the hypothesis that obligatory components of the ERP that are sensitive to physical properties of sounds can influence measurements of MMN. Intensity modulation of model responses is clearly observed in Figure 2, Figure 3, and Figure 5. In particular, hidden unit activations within the latency ranges of P50 and N1 peaks were significantly influenced by stimulus intensity level. Interestingly, although simulated 70 dB and 90 dB deviant stimuli produced model outputs with opposite polarity amplitude shifts over the 125 to 250 ms measurement window associated with MMN, units that peaked within this latency range were not significantly affected by input stimulus level, suggesting that the activity of earlier components spreads over to influence ERP amplitudes during this range. This aligns with some of the arguments put forth by May (2021), namely that MMN is mistakenly interpreted as a separate component, and actually arises due to amplitude and latency modulation of neural generators that ordinarily precede the latency of MMN measurement. These findings are also consistent with a previous analysis of this dataset, which found that rising and falling sound level transitions produced opposite polarity changes in ERP amplitudes over the measurement window of MMN (O’Reilly, 2021b). However, this presents an apparent contradiction to the predictive coding account of auditory processing, which views MMN as evidence for a specific, dissociable component of the ERP that reflects prediction-error signalling (Carbajal and Malmierca, 2018; Garrido et al., 2009; Näätänen et al., 2005; Shröger, 2007).

While it might be difficult to square these, there is at least one perspective from which to interpret intensity dependence of the ERP within the context of predictive coding. Recently some have postulated the existence of different positive and negative prediction error units that encode unexpected stimuli that are above or below expectations, respectively (Hertäg and Clopath, 2022; Keller and Mrsic-Flogel, 2018). Evidence in support of this hypothesis comes from foundational studies of reward prediction-error (Schultz and Dickinson, 2000). With reference to the current findings, differences between model responses to 90 dB and 80 dB stimuli could be described as a positive prediction-error unit signal. Conversely, differences between 70 dB and 80 dB responses could be defined as a negative prediction-error unit signal. However, this interpretation does not intrinsically account for differences in polarity between resulting mismatch responses, and it is unclear how the abstract notion of over- and under-prediction of sensory information relates to the encoding of complex spectral-temporal properties of sounds. Moreover, while an interesting theory, the benefits of adopting this framework to describe the current data are not obvious, given the relative paucity of evidence for real prediction and prediction-error neurons (Heilbron and Chait, 2018; May, 2021), in contrast with intensity-modulated sensory neuronal activity, which is a comparatively well-established phenomenon (Bajo and King, 2012; Duque et al., 2016; Hart et al., 2002; Langers et al., 2007; Röhl and Uppenkamp, 2012).

It has also been pointed out that control sequences designed to verify interpretations of MMN may overlook some of aspects of sensory processing that can influence responses to control stimuli (O’Reilly and O’Reilly, 2021). For example, the roving oddball sequence induces repetition suppression and may also reflect asymmetric sensitivity of the auditory response to the physical parameter(s) under investigation (O’Reilly, 2021a). Additionally, contrast gain control (Lohse et al., 2020; Rabinowitz et al., 2011) presumably modulates responses to different tones presented in many-standards sequences, which include a range of physically different stimuli, thereby confounding analysis between responses to deviant and control stimuli (Parras et al., 2017; SanMiguel et al., 2021). Furthermore, other sources of interference with putative MMN measurements may emerge as unintended consequences of subject behaviour during passive auditory oddball experiments. For instance, two common approaches are to instruct the subject to watch a silent film or read a book, which present different sets of cues that could unintentionally modulate the auditory response in different respects (Biau et al., 2021; Rayner and Clifton, 2009; Zoefel, 2021). As such, demonstrating that MMN reflects a specific prediction-error signal is challenging, and the balance of evidence from the current study leans decidedly towards the hypothesis that intensity MMN reflects modulation of obligatory sensory components of the ERP. This highlights the inherent difficulties involved in disentangling hypothetical expectation and prediction components from intrinsic sensory components measured from scalp-recorded potentials.

The hierarchical RNN potentially offers a valuable model for studying the computational principles underlying ERP generation, and more broadly in cognitive neuroscience research (Barak, 2017; Yang and Molano-Mazón, 2021). Figure 2 and Figure 3 illustrate why the hidden units of the model may be considered to behave loosely analogously to biological neural sources which give rise to scalp-recorded ERPs. These figures demonstrate that the model produces sequences of internal states that increase in complexity from superficial to deep layers, as information travels through the network, which presents a parallel with cortical sensory processing. The results from PCA in Figure 4 suggest that (i) model responses are separable based on stimulus level (70 dB vs. 80 dB vs. 90 dB) and stimulus type (80 dB standard vs. 70 dB and 90 dB deviants), (ii) deeper layers (3 and 4) explain more variance than superficial layers (1 and 2), and (iii) hidden unit temporal response categorisations are reflected in principal component space. Importantly, the simulated 90 dB stimulus provokes a reasonable-looking simulated ERP response from the model, comparable with previous findings (Hagenmuller et al., 2016; Morrison et al., 2019; O’Reilly, 2021b). Despite this apparent realism, we cannot ascertain whether the model outputs truly match the grand-average ERP produced by a 90 dB stimulus, given that this stimulus condition was not included in the original experiment (Kappenman et al., 2021). Moreover, it should be recognised that the model is subject to the same limitations of conventional grand-average ERP analysis (Luck, 2014), since it was not trained to replicate individual subject ERPs, and that terms ‘shallow’ and ‘deep’ used in the context of this model imply no relation to cortical laminae.

## 5. Conclusions

This article has presented a technique for modelling and analysing human event-related potentials recorded during passive auditory oddball stimulation. The multi-layered RNN effectively generates synthetic ERP waveforms by the combined actions of its hidden units, which offer an analogy to biological neural sources. Although subject to some of the same limitations as conventional grandaverage ERP analysis and source-based techniques, this approach may nevertheless offer considerable value in trying to understand the computational principles underlying ERP generation. The results of this modelling study support the view that sound intensity modulates ERP amplitudes during the latency range of MMN. This may be interpreted as loudness dependence of sensory ERP components that cause amplitude changes that spill over into the measurement window for MMN, which agrees with converging electrophysiological findings in schizophrenia patients (Hagenmuller et al., 2016; Todd et al., 2008; Wang et al., 2022). Alternatively, this could be interpreted in terms of a recent rensition of the predictive coding framework (Hertäg and Clopath, 2022; Keller and Mrsic-Flogel, 2018), whereby opposite polarity amplitude changes in response to quieter and louder deviant stimuli may be taken to reflect examples of negative (i.e. below expectations) and positive (i.e. above expectations) prediction errors, respectively. To move forward productively, these interpretations ought to be verified in future experimental and computational studies designed to distinguish between the neurophysiological mechanisms responsible, on the one hand, for loudness dependence, and on the other, for predictive modelling and prediction-error signalling.

## Acknowledgements

I am sincerely grateful for researchers who are practicing openness and transparency in research, and specifically for Emily S. Kappenman, Jaclyn L. Farrens, Wendy Zhang, Andrew X. Stewart, and Steven J. Luck for making the ERP CORE resources freely available. The model developed in this study was trained using a computer system and graphics processing unit purchased with support from the Research Institute of Rangsit University [grant number 90/2561].

## Data availability

The raw data used in this study is available to download from https://doi.org/10.18115/D5JW4R, and the code and model can be accessed from *[repository will be shared prior to publication].*

## Ethical review

The experimental work performed to obtain the data used in this study was previously approved by the ethics committee of the originating researchers.

## Author Contributions (CRediT Statement)

Jamie A. O’Reilly: Conceptualization, Methodology, Software, Formal analysis, Data Curation, Writing - Original Draft, Writing - Review & Editing, Visualization.

